# Insights on *Chamelea gallina* growth dynamics from the Holocene climate optimum in the Northern Adriatic Sea (Italy)

**DOI:** 10.1101/2025.04.09.647857

**Authors:** Alessandro Cheli, Arianna Mancuso, Fiorella Prada, Alexis Rojas, Giuseppe Falini, Stefano Goffredo, Daniele Scarponi

## Abstract

The fossil record documents ecological responses to past climate transitions, providing insights into future scenarios of marine ecosystems and taxa under climate change. In this study, we compared shell growth dynamics, specifically linear extension and net calcification rates, of the bivalve *Chamelea gallina* between Northern Adriatic Sea (NAS) assemblages from the Holocene Climate Optimum (HCO, 9-5 kyr cal. B.P.) and today. This species is a valuable economic resource, currently threatened by climate warming and numerous anthropogenic stressors. During the HCO, regional sea surface temperatures were warmer than today, making it a potential analog for exploring ecological responses to increasing seawater temperatures predicted in the coming decades. By combining standard aging methods with reconstructed sea surface temperatures, we observed a significant reduction in linear extension and net calcification rates in warmer HCO assemblages. During the HCO, immature *C. gallina* specimens developed a denser shell, especially during their early growth stages, at the expense of a linear extension rate, which was lower than modern specimens. This resulted in an average delay of 3-4 months in reaching sexual maturity, which is currently reached after ~ 13-14 months or at a length of ~18 mm. Several environmental factors are probably responsible for these differences between fossil and modern assemblages. Temperature indirectly impacted the geomorphologic evolution of the NAS over the Early-Mid Holocene, shaping the outlet structure of the Po and other NAS rivers. This, in turn, affected environmental factors such as nutrient load, seawater transparency, salinity, and phytoplankton. In addition, recent anthropogenically-derived decreases in natural NAS predators of infaunal bivalves may have reduced the natural predation pressure on modern *C. gallina*, thus favoring those populations characterized by faster linear extension rates (at the expense of a higher shell density), especially during the initial stages of life, hence facilitating a quicker attainment of sexual maturity.

## 1. Introduction

Studies on the capacity of marine organisms to acclimatize or adapt to temperature changes have increased during the past few decades as the severity of climate warming has been recognized (Hofmann et al., 2011; Kroeker et al., 2016; Ninokawa et al., n.d.; Wahl et al., 2015; Wootton et al., 2008; Wootton and Pfister, 2012). Some coastal areas have experienced changes in mean seawater temperature, in some cases equal to or greater than the conditions originally predicted by future ocean scenarios (Hofmann et al., 2011; Kroeker et al., 2016; Wahl et al., 2015; Wootton et al., 2008; Wootton and Pfister, 2012).

A good understanding of long-term biotic dynamics in the context of major environmental shifts is required to reduce the uncertainty regarding organismal responses to ongoing climate (and derived environmental) changes. While laboratory studies enable the isolation of a specific stressor, field-based ecological studies are necessary to better understand the combined effects of multiple stressors in nature. Nevertheless, detecting long-term population dynamics in field-based ecological studies can be time-prohibitive. Temporally well-resolved (i.e., low time-averaged) fossil-rich assemblages retrieved in sedimentary successions might record ecological responses to past climatic-driven environmental shifts, far beyond the limited timescales of direct ecological monitoring, typically restricted to the most recent decades (Huntley *et al*., 2014; Kidwell, 2015; Pfister *et al*., 2016; Scarponi *et al*., 2017; Tomašových *et al*., 2020; Finnegan *et al*., 2024; Nawrot *et al*., 2024). Given their extensive fossil record, marine calcifying organisms are unique recorders of past environmental changes (Bemis et al., 1998; Chauvaud et al., 2005; Posenato et al., 2022; Rhoads and Lutz, 1980; Schöne and Gillikin, 2013; Vihtakari et al., 2017). In particular, bivalve shells record historical data due to their seasonal deposition of carbonate material, retaining high-resolution temporal records of the ambient physical and chemical conditions during growth (Klein et al., 1996; Purroy et al., 2018; Schöne et al., 2003).

The venus clam *Chamelea gallina* has been increasingly used as a model organism for evaluating adaptation responses to near-future environmental changes (Gallagher and Albano, 2023; Moschino et al., 2023). This species is abundant in shoreface settings at depths ranging from 0 to 12 m up to 1–2 nautical miles in the Mediterranean Sea (Froglia, 1989). It is particularly abundant in the western Adriatic Sea (Mediterranean Geographical Subareas-GSA 17), where the massive outflow of the Po (along with that of other minor rivers) and the anti-clock-wise currents in the NAS provide, along the Italian coasts (and particularly around and south of the Po Delta), abundant nutrients, particles, and organic matter (Orban et al., 2007). Moreover, *C. gallina* represents an economically and ecologically important species in this basin, and its harvesting in the GSA 17 accounts for over 98% of ‘venus clams’ production at the national level (Scarcella and Cabanelas, 2016). Previous studies have mainly focused on population dynamics, shell growth, and composition of this species in populations living along the western Adriatic coasts (Bargione et al., 2020; Carlucci et al., 2024; Gizzi et al., 2016; Grazioli et al., 2022; Mancuso et al., 2019). It has been observed that the shell properties of *C. gallina* populations are influenced by environmental changes, with variations in shell density, thickness, and growth primarily related to varying water temperatures and secondarily to salinity and food availability along a spatial gradient (Gizzi et al., 2016; Mancuso et al., 2019). Along the latitudinal gradient in the Adriatic Sea, shells of the warmest populations were characterized by lighter, thinner, more porous, and fragile shells (Gizzi et al., 2016).

The long-term dynamic of *C. gallina* shell properties and its microstructural features (i.e., bulk density, microporosity, micro density) were previously investigated in response to past climate-driven environmental changes on a millennial temporal scale by comparing fossil (~7.6, ~5.9 and ~2.6 ky BP) and modern assemblages of the western NAS (Cheli et al., 2021). Specifically, modern assemblages displayed less dense and more porous shells than fossil ones, likely due to lower present-day aragonite saturation states (Cheli et al., 2021). The current study builds upon these findings to evaluate linear extension and net calcification rates of *C. gallina* in relation to Holocene climate-driven environmental changes that occurred in the NAS south of the Po Delta. Four out of five shoreface-related *C. gallina* assemblages previously investigated by Cheli and collaborators were considered here: two from the present-day NAS (namely MGO and MCE; Fig. S1) and the two from the HCO (i.e., CO1 and CO2, respectively dated 7.6 ± 0.1 ky B.P., and 5.9 ± 0.1 ky B.P.; Fig. S1), when regional sea surface temperatures were higher than today. Indeed, reconstructed SST showed a difference of approximately 1.5 °C between the lower HCO (i.e., CO1, 18.6°C) and present-day (MCE, 17.3 and MGO, 17.1°C; Table 1), thus representing a possible analog for the sea temperatures predicted for the next decades. This study addresses the knowledge gap concerning the natural range of variability of *C. gallina* at temporal scales beyond those typically considered in ecological monitoring, providing valuable insights into the potential responses of *C. gallina* to near-future global warming.

**Table 1.**
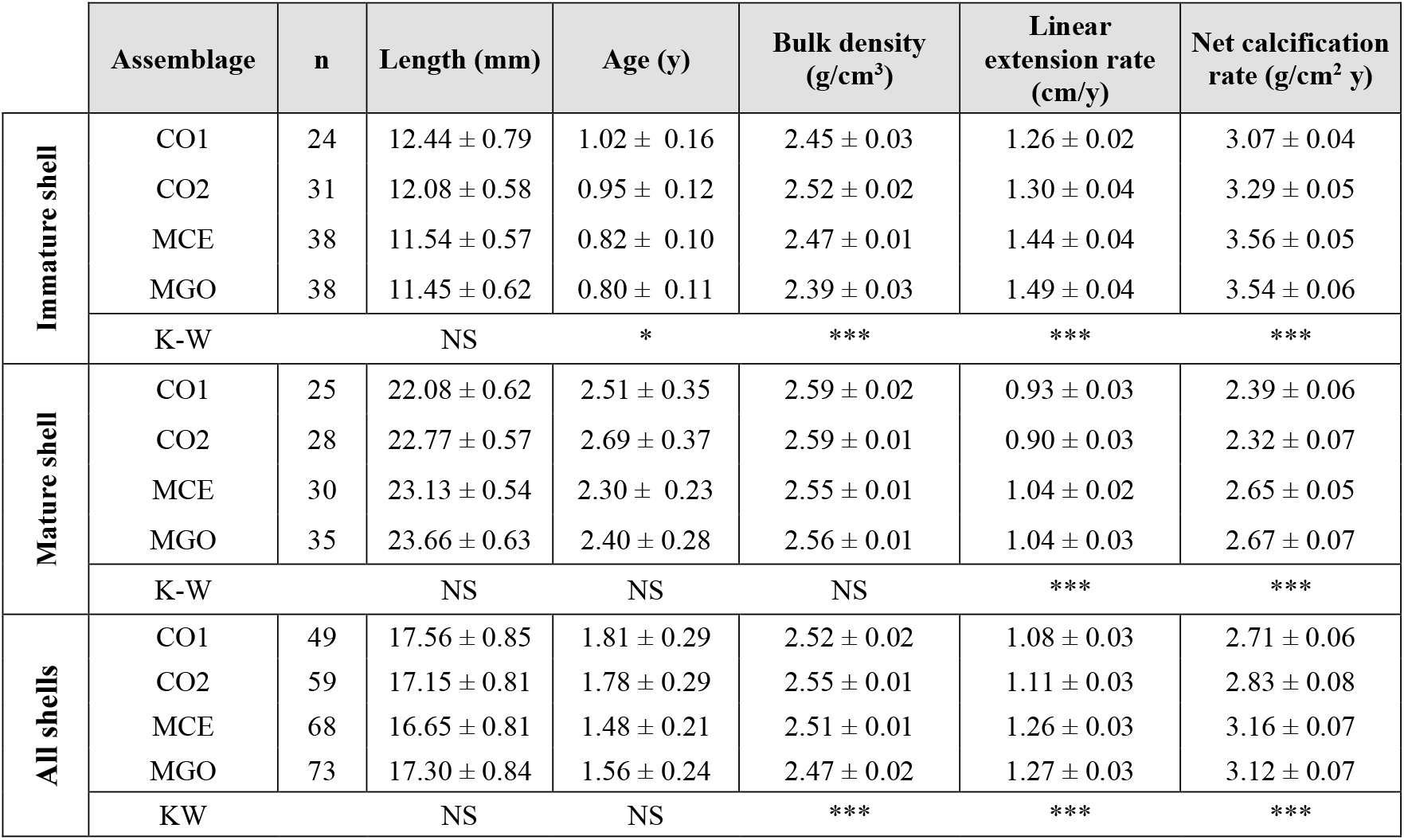
Calibrated radiocarbon age, sea surface temperature (SST), and Von Bertalanffy growth function (VBGF) parameters. Values for each assemblage in chronological and temperature order. Radiocarbon measurements are reported in years ‘before present’ (B.P.). L_inf_ and k were estimated from the generalized VBG function by pooling the data for external and internal rings. n = a number of collected specimens. K-W = Kruskal-Wallis rank test; NS not significant, *** p < 0.001. Assemblage labels, number of right valves examined (n), calibrated ages (^14^C Age), and Sea Surface Temperature (SST) are after Cheli et al. (2021).

## 2. Materials and methods

### 2.1 Study area and sampling

The NAS is a semi-enclosed, shallow basin characterized by a wide shelf with a low topographic gradient of 0.02° and an average depth of 35 m (Poulain, 2001). It extends approximately 350 km southward from the Gulf of Trieste and is bordered westward by Italy and eastward by the Balkan Peninsula. The coasts along the Adriatic western (Italian) sector are primarily sandy and strongly influenced by the Po River outflow, which affects the water circulation in the northern Adriatic Sea and plays a fundamental role in the bio-geochemical processes of the basin (Marini et al., 2008). The Po is the largest Italian river, and currently, it supplies over 50 % of freshwater input to the NAS (Degobbis et al., 1986) and about 20 % of the total river discharge in the Mediterranean Sea (Russo and Artegiani, 1996). However, the Adriatic sedimentary succession records major changes in the basin configuration in relation to the last glacial-interglacial transition (Amorosi et al., 1999; Gamberi et al., 2020). During the early Holocene Climate Optimum (HCO, 9–7 ky B.P.), the north-western Adriatic coast was characterized by estuary systems bounded seaward by a series of lagoons that limited riverine plumes into the Adriatic Sea (Amorosi et al., 2019). By contrast, during the last part of the HCO (i.e., between 7.0 to 5 ky B.P.), the area transitioned firstly to a wave-dominated and, after 2.0 ky B.P., to a river-dominated deltaic system (Amorosi et al., 2019). The last geomorphologic configurations led to the progressively increasing influence of the riverine processes that control present-day coastal dynamics and storage-release of sediments and nutrients. The enhanced freshwater discharge in the nearshore area, especially during the last 2.0 ky B.P., resulted in a strong progradation and the upbuilding of the modern Po Delta (Correggiari et al., 2005).

Our analyses focused on four *C. gallina* assemblages reported by Cheli et al., 2021 and grouped in two specific time intervals: 1) the HCO (at ~7.6 and ~5.9 kyr cal. B.P.) documented in subsurface cores and 2) the present day represented by the surficial dead assemblages of the NAS (Fig. S1). Details on age determination methods, sampling design, specimen collection and preparation, and estimate of environmental parameters can be found in Cheli et al., 2021, with a brief description provided in the supplementary material.

### 2.2 Age determination of *C. gallina* specimens

The specimen’s age was estimated in a subsample of 30 shells of different sizes in each assemblage using three aging methods: shell surface growth rings, shell internal rings, and (partly) stable oxygen isotope composition (Fig. 1). The growth curves were obtained by counting the total number of visible external and internal rings in each examined valve and then fitted with the von Bertalanffy growth function (VBGF): *L*_*t*_ = *L*_*inf*_[1 − *e*^−*k*(*t*)^]

**Figure 1.**
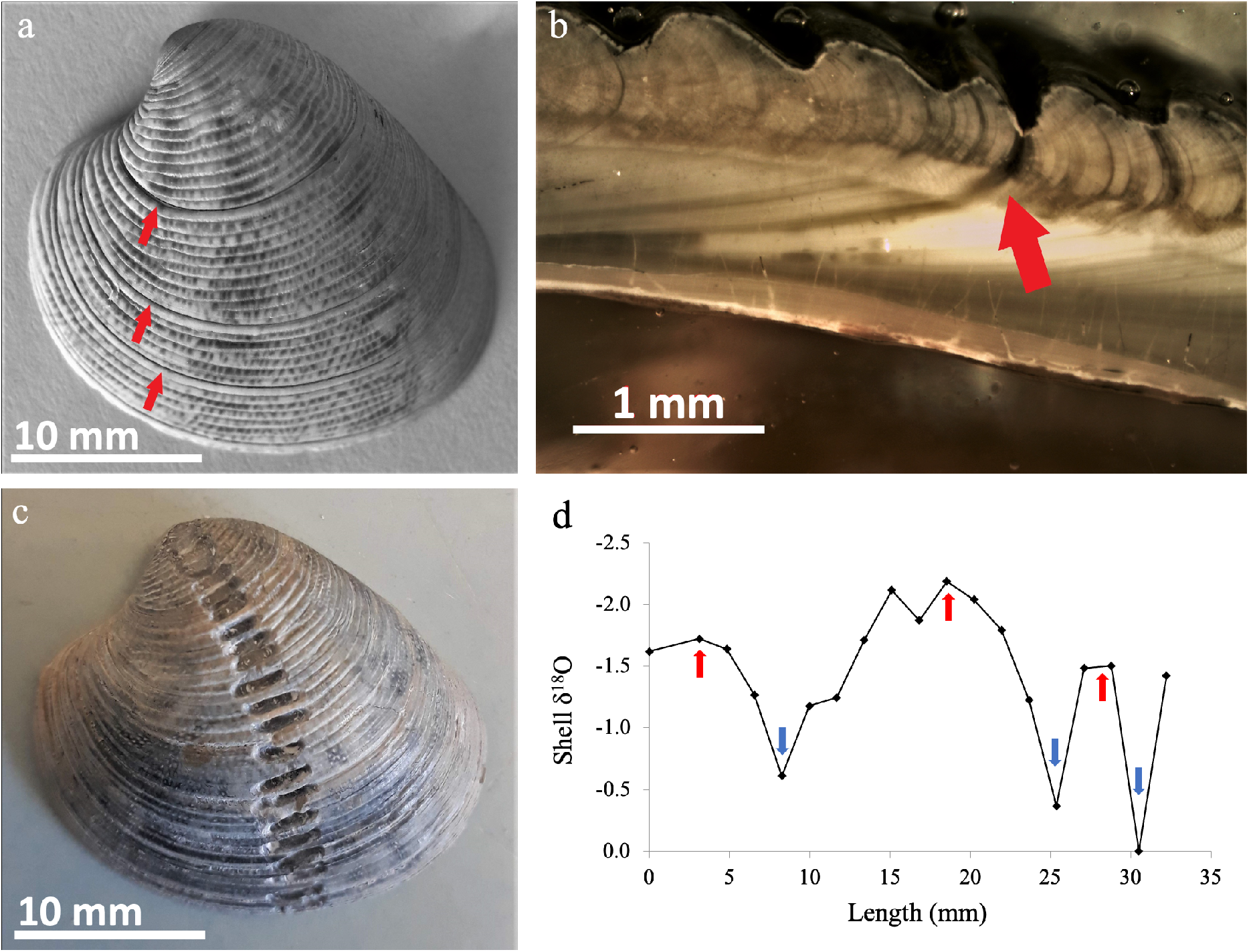
Shell ageing methods. (a) external growth rings on the surface of *C. gallina* after shell scanning; (b) internal growth rings in the shell section; (c) samples of *C. gallina* with drilling “spots” from the umbo to the ventral edge to determine δ^18^O values along the shell growth axis; d) sinusoidal sequence of lower (summer, red arrows) and higher (winter, blue arrows) δ^18^O values recorded by the shells.

L_t_ is the individual length (the maximum distance on the anterior-posterior axis) at age t, L_inf_ is the asymptotic length (the maximum expected length in the population), k is the growth constant, and t is the specimen’s age. Hence, two growth curves for each specimen belonging to a studied assemblage were produced, and a chi-square test of maximum likelihood ratios was used to examine the significance of differences in growth functions between the two aging methods. If differences are present between VBGF curves from valve external and internal ring counting methods, the growth curve will be based on internal ring counting, which is considered more reliable (Bargione et al., 2020). Whereas, if no differences are revealed between VBGF curves from the two-counting methods, a generalized growth curve will be constructed for specimens of each assemblage by merging the growth curves from both methods (Table 1; Fig. 2). The resulting generalized VBGF curves were used to extrapolate the age in all the other shells by applying the inverse of the generalized VBGF:

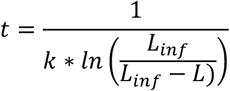

**Figure 2.**
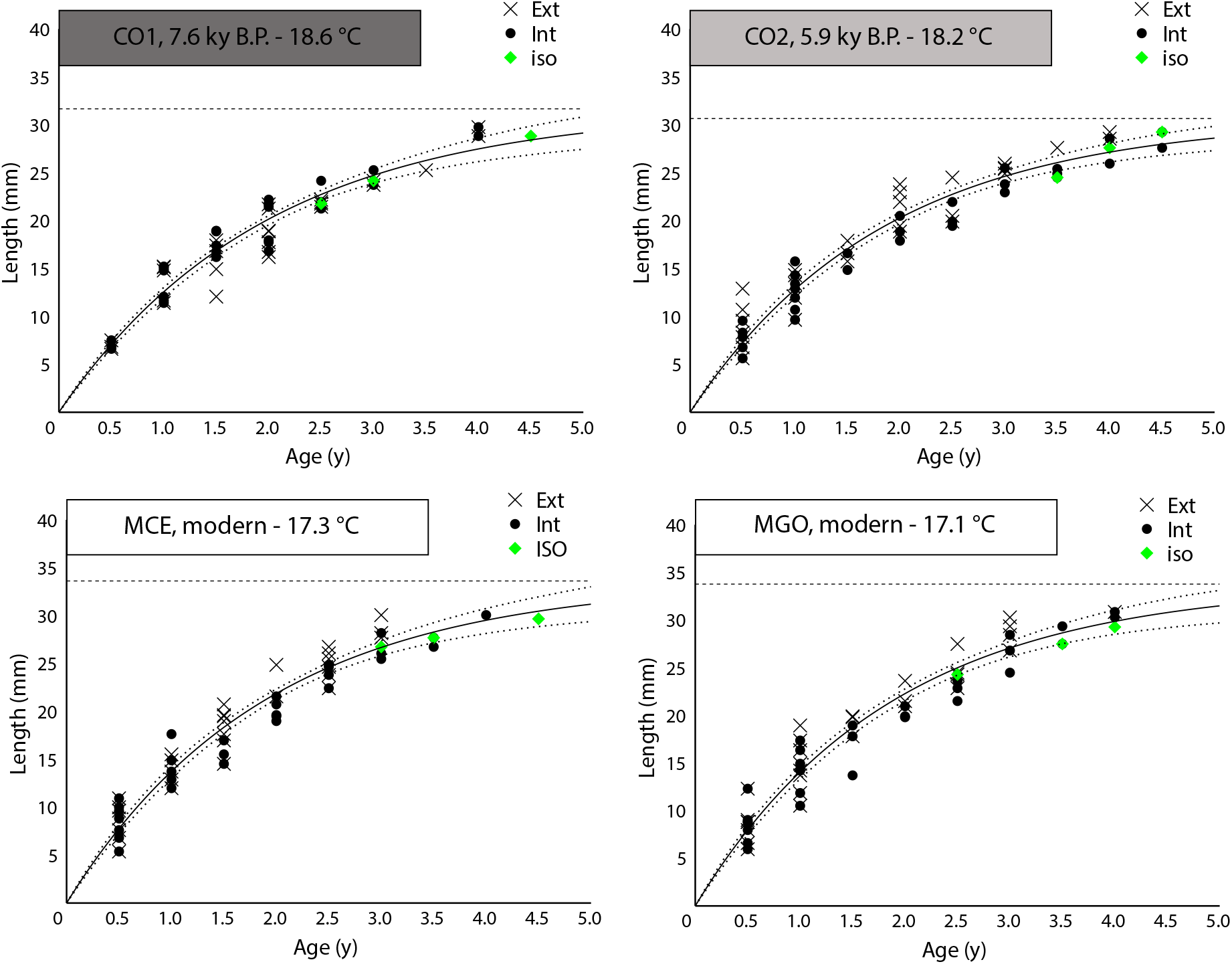
Von Bertalanffy growth curves. The generalized age-length von Bertalanffy growth curve for each assemblage was obtained by merging data from the two aging methods (counting external and internal rings). Green diamonds represent the ages obtained from the δ^18^O profiles along the shell growth axis and serve as validation to the counting aging methods by fitting the curves.

To validate the data from the two aging methods and the generalized VBGF curves, δ^18^O measurements on three valves per assemblage were carried out at the Stable Isotope Biogeochemistry Laboratory, Department of Earth, Environmental, and Planetary Sciences, Washington University in St Louis (MO-USA). δ^18^O measurements were carried out on “spot” samples collected from the prismatic and cross-lamellar layers and drilled sequentially along the shell growth direction by a single diamond drill. The age-length estimation resulting from counting lighter δ^18^O (summer) and heavier δ^18^O (winter) peaks were plotted alongside the age-length key from the two aging methods (Fig. 2).

### 2.3 Determination of linear extension and net calcification parameters

In *C. gallina* specimens, linear extension rates were obtained from the valve length/age ratio (cm y^−1^), while the net calcification rate (mass of CaCO_3_ deposited per year per unit area, g cm^2^ y^−1^) was calculated for each valve by applying the formula: bulk density (g cm^−3^) x linear extension rate (cm y^−1^). The bulk density of each valve examined in our study was sourced from the literature (see Table 2 in Cheli et al., 2021). Correlation analyses between linear extension rates, net calcification rates, and SST were performed to test for any significant relationship developed over geological time as a function of temperature-driven environmental changes. Differences in the growth of *C. gallina* specimens were also examined in relation to sexual maturity, which is reached at around 18 mm in modern times (Grazioli et al., 2022), to account for potential variations in the biomineralization process during different stages of the bivalve’s life cycle.

### 2.4. Statistical comparison of shell parameters among assemblages

Since assumptions for parametric statistics were not fulfilled, the non-parametric Kruskal-Wallis equality-of-populations rank test was used to test for differences in shell parameters among assemblages, followed by Dunn’s post hoc tests. Mann–Whitney U tests were applied to determine the differences between ages obtained from external and internal growth rings within each assemblage. For each assemblage, the age-length functions were fitted with the VBGF using a non-linear model, which estimated the VBGF parameters through bootstrapping. Kimura’s likelihood test was then used to compare VBG functions among assemblages. Regarding age in each assemblage, logarithmic regression models were fitted to shell bulk density, while linear models were fitted to linear extension and net calcification rates. Spearman’s rank correlation coefficient (rho) was used to evaluate trends between shell growth parameters and sea surface temperature. All statistical analyses were computed using RStudio software (RStudio Team, 2022).

## 3 Results

### 3.1 Specimen age-length von Bertalanffy growth curve

In all examined assemblages, there is no significant difference in the extrapolated ages of fossil and modern *C. gallina* specimens between estimations based on external and internal rings (Mann-Whitney *U* test, p > 0.05). Moreover, Kimura’s likelihood ratio tests indicated no significant differences between the growth curves obtained from the external and internal rings. Therefore, a generalized VBGF curve was obtained for each assemblage using the combined data (Fig. 2). Measurements of stable oxygen isotopes (δ^18^O) validated the extrapolated ages from the aging methods by fitting the VBGF curves (Fig. 2).

Differences in the observed values for the estimated mean asymptotic specimen length (L_inf_) (anterior-posterior) and the growth constant (k) in the targeted assemblages were not statistically significant (Kruskal-Wallis test, df=3 and p > 0.05; Table 1). Shell length and estimated life span were comparable among assemblages for mature and all shells (Kruskal-Wallis test, df = 3 and p > 0.05; Table 2). The age at the onset of sexual maturity resulted higher in the warmer HCO assemblage (i.e., +28% in CO1) compared to modern ones, thus showing a delay in reaching sexual maturity in the former (Kruskal-Wallis test, df = 3 and p < 0.05; Table 2).

### 3.2 Specimen linear extension and net calcification parameters

In each assemblage, the examined shell growth parameters (i.e., linear extension and net calcification rates) were negatively correlated with shell age (Fig. 3). Linear extension and net calcification rates differed significantly among assemblages (Table 2, Fig. 4). Bulk density was significantly different for immature specimens and hence for the whole dataset among assemblages (Table 2, Fig. 4). The variation of the two above-mentioned growth parameters was then analyzed in relation to temperature-driven changes between the HCO and the present day. Linear extension rate and net calcification rate showed a marginally significant negative correlation with SST in the whole dataset (rS = −0.311 and −0.292, respectively, p<0.001; Fig. 4). Correlations with the SST were also performed separately for immature and mature specimens, with both sub-datasets exhibiting a significant negative correlation for linear extension and net calcification rates (Fig. 4). The two assemblages positioned at the highest and lowest temperature range over time (i.e., CO1–18.6°C and MGO– 17.1°C) displayed stronger variations in immature specimens, with a 2.3 mm (~18%) per year difference in mean linear extension rate and a 0.47 g/cm^2^ (~15%) per year difference in mean calcification rate and (Table 2, Fig. 4).

**Figure 3.**
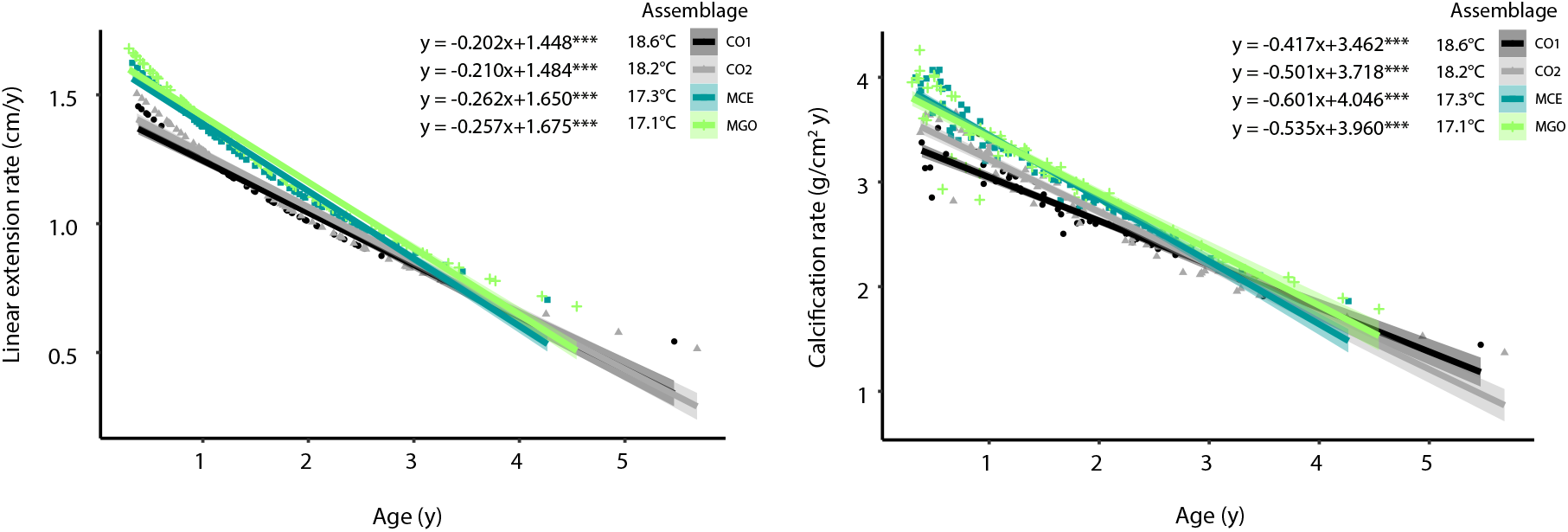
Correlation of the growth parameters with shell age in the four different assemblages. The regression line and 95% confidence interval for predictions from a linear model (‘lm’) are visualized using the geom_smooth function of the R ggplot2 package. *** p<0.001.

**Figure 4.**
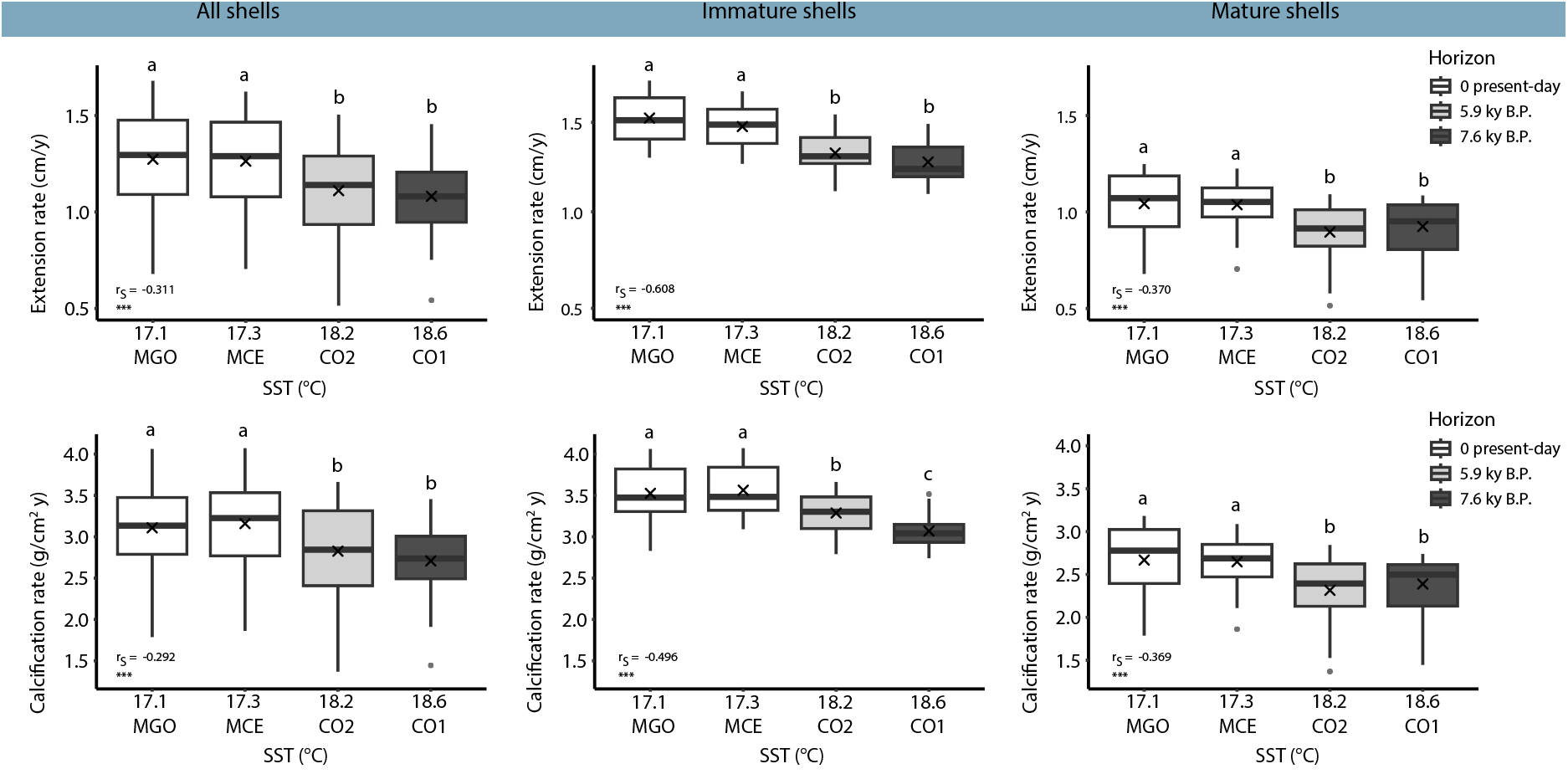
Variation of shell growth parameters in relation to different SST (Sea Surface Temperature). Three groups were considered: correlations in all shells, correlations in immature shells (<18 mm), and correlations in mature shells (>18 mm). The box colors represent the age of the assemblages, ranging from the oldest fossil horizon (dark grey boxes) to the present-day horizon (white boxes). The boxes indicate the 25^th^ and 75^th^ percentiles, the lines within the boxes mark the medians, and the crosses mark the mean values. Black points represent outliers. Different letters indicate statistical differences among assemblages (p < 0.05; the number of clams measured for each assemblage is reported in Table 1). rS, Spearman’s rho coefficient. ***p < 0.001, NS, not significant.

## 4. Discussion

The radiocarbon analyses of *C. gallina* specimens revealed that fossil and modern dead assemblages have a comparable time averaging (i.e., the cumulative amount of time in which individuals forming an assemblage have lived). Radiocarbon dating of seven *C. gallina* specimens from a nearby shoreface dead assemblage showed an interquartile range (IQR) of 50 years (Fitzgerald et al., 2024). Along the same line, the radiocarbon dating of eleven *C. gallina* specimens from sample CO1 resulted in an IQR of 85 years (Table S1). This suggests that the fossil and modern dead assemblages have accumulated over similar time spans (decades-century). As a result, the trends in biomineralization observed in the analyzed assemblages reflect long-term, mediated responses to environmental changes and buffer the results from possible short-term or uncommon environmental perturbations that could have affected a specific cohort of the *C. gallina* population.

### 4.1 Estimates of maximum length and growth parameters in *C. gallina*

Despite the variation in temperature between the Holocene climate optimum (HCO) and present-day conditions, the maximum expected length of the population (L_inf_) and the rate at which it is approached (k) do not show significant changes between fossil and modern assemblages (Fig. 2; Table 1). The estimated mean asymptotic shell lengths (L_inf_) for fossil and modern assemblages were lower than those reported in previous studies for modern living populations of *C. gallina* from the Adriatic nearshore settings (Arneri, E and Froglia, C and Polenta, R and Antolini, 1997; Bargione et al., 2020; Deval, 2001; Gaspar, 2004; Mancuso et al., 2019; Ramón and Richardson, 1992). However, considering the lower confidence interval for L_inf_ observed in the sites around the Po delta, the modern shells from this study were comparable with the modern ones from Mancuso et al., 2019. The observed values of von Bertalanffy growth constants (k) for all assemblages were consistent with nearly almost all previous data obtained in the Adriatic Sea (Polenta R., 1993; Arneri, E and Froglia, C and Polenta, R and Antolini, 1997; Mancuso et al., 2019). A lower L_inf_, but the same k suggests that while individuals in one population will not grow as large as those in another, they grow at a comparable rate during their life span. This could be influenced by a combination of environmental, genetic, and ecological factors that limit the maximum achievable size without affecting the intrinsic growth rate. On a decadal scale, the specimens of the present study might have lived in conditions of lower nutrient availability and higher competition for resources (i.e., higher density; Peterson & Beal, 1989), which affected the L_inf_ value but not their intrinsic growth rate. These variations in the von Bertalanffy growth equation parameters describing assemblages in space and time highlight the high plasticity of this species in adapting to changing environments.

### 4.2 Microstructural parameters affecting the growth of *C. gallina*

At a broad scale (specimen life span), our results support the general physiologic decrease in linear extension rates (and also net calcification rates) as a function of specimen age observed in mollusks and other organisms (Caroselli et al., 2016; Mancuso et al., 2019; Sebens, 1987). For example, in living specimens of *C. gallina* retrieved along the targeted sector of the NAS (Mancuso et al., 2019), the linear extension rate of *C. gallina* decreased with increasing age, highlighting fast growth in the first year of life, with a reduction of around 20% in the second year and around 40% in the third year. Hence, the power-shaped relation between growth parameters and age is confirmed in dead assemblages from modern NAS and HCO samples (Figure 2). This suggests that the overall growth dynamics did not change even when temperatures were higher (1.5 °C) than present-day and comparable to temperature estimates forecasted for the NAS in the next few decades (i.e., before the end of this century (Branković et al., 2013).

Significant differences in growth patterns emerge when data are analyzed at the subgroup level, especially during the juvenile phase. Specimens from fossil horizons, which lived in warmer coastal settings of the NAS characterized by a reduced fluvial influence due to the estuarine configuration of the Po and other rivers, exhibited denser shells with lower extension and net calcification rates than modern Adriatic specimens located south of the Po Delta (Fig. S1; Table 2). Specifically, linear extension rates were depressed during the HCO with respect to present-day assemblages. From a thermodynamic point of view, higher temperatures generally promote CaCO_3_ deposition by decreasing the energetic costs of shell formation with an increasing aragonite saturation state (Clarke, 1993). Our data suggest a possibly higher saturation state scenario in the NAS during the Holocene; such a scenario favors the precipitation of a denser shell in warmer seawater conditions. Similarly, the net calcification rates of *C. gallina* during the HCO are significantly depressed during the juvenile and mature stages of life. So, at present, juvenile clams in the investigated area may be allocating more energy towards faster growth (linear extension rate) at the expense of skeletal density to reach size at sexual maturity. Comparatively, juvenile specimens of *C. gallina* from the HCO reached maturity with a delay of around three months, with respect to the modern juvenile specimens that attained maturity after thirteen-fourteen months in the modern NAS (Mancuso et al., 2019). This delay could represent an important factor for the near-future management of this economically valuable resource. Higher temperature usually coincides with greater and faster gonadal development in bivalves, although this is mainly due to a greater amount of food ingested (Delgado and Pérez Camacho, 2007). However, a previous study conducted on the clam *Ruditapes philippinarum* found that a high temperature of around 18°C is associated with a low ingestion rate, a situation of negative energy balance arises, leading to a slow rate of gonadal development that takes place at the cost of the animal’s energy reserves (Delgado and Pérez Camacho, 2007). This behavior could explain the growth pattern of *C. gallina* in HCO due to physical factors dependent on a warmer sea (influencing the aragonite saturation state), combined with lower nutrient run-off from the Po River (with respect to present-day).

It is also interesting to note that within the same study area, we have been assisting in the last centuries to a generally decreasing trend in predator-prey interactions, especially between shell-drilling gastropods and their molluscan prey as testified recently by Zuschin and colleagues on slightly deeper (prodelta) environments from the same targeted area (Zuschin et al., 2024). Here, an unprecedented (relative to the Holocene) increase in dimension and population of *Varicorbula gibba* (another infaunal bivalve) was observed, and it was hypothesized to be driven by ecological release from predation (Zuschin et al., 2024). Thus, the recent removal of predators (naticids) feeding also on *C. gallina* and other mollusks (La Perna, 1992) might have released *C. gallina* and favored those populations characterized by faster linear extension rate to reach sexual maturity faster. Conversely, denser conchiolin layers retrieved in *C. gallina* specimens from HCO can provide a buffer against drilling predators (Kardon, 1998) and could also lead to a selective advantage, especially during their juvenile stages, where predation is more frequent (Zuschin *et al*., 2024). However, this resulted in an overall decrease of linear extension rates in *C. gallina* that lived in a warmer Adriatic Sea, thus taking more time to reach the maturity size.

### 4.3 The response of *C. gallina* to warming

Based on the current knowledge of the main limiting factors acting on this species’ growth and the results obtained by *C. gallina* cohorts that lived mainly during the early HCO, we can discuss the principal factors affecting *C. gallina* populations in the near future, that is seawater temperature and nutrients.

#### Low tolerance to high seawater temperatures

Previous studies highlighted the relatively low tolerance of *C. gallina* to high temperatures compared with other bivalve species, showing a great influence on the overall physiological responses and heavy stress conditions when exposed to high temperatures, demonstrating that temperature could be a tolerance limit for this species (Matozzo et al., 2012; Monari et al., 2007; Moschino and Marin, 2006). Indeed, specimens of *C. gallina* exposed to high summer temperatures (28 °C) showed reduced energy absorption and increased energy expenditure via respiration, negatively affecting the energy balance (Moschino and Marin, 2006) and probably growth, as found during the summer season (temperature > 27 °C) in specimens from the eastern coast of Spain (Ramón and Richardson, 1992). Moreover, oxygen depletion due to high temperatures may harm the clam’s physiological performance, as observed in the bivalve *Ruditapes decussatus* (Sobral and Widdows, 1997). This study evaluated the growth patterns of *C. gallina* in a warmer context than present-day conditions (~ 1.5 °C); this would imply for HCO longer time, especially during summer seasons where temperatures could have intercepted the threshold of suboptimal growth, thus depressing linear extension rates of *C. gallina* during the summer season. Following (Branković et al., 2013), the mid-twenty-first century climate scenario for the northeast Adriatic region depicts an increase in sea surface temperatures reaching + 3.5 °C in summer and early autumn (Branković et al., 2013). Thus, *C. gallina* growth should be strongly negatively affected by climate warming during the summer, given that present-day summer average SST temperatures (June-September) in the targeted portion of the Adriatic are 25-26 °C. Therefore, *C. gallina* could show a strong contraction (> 90%) of its distributional range well before the end of the century (Gallagher and Albano, 2023).

#### Nutrient supply

The growth of *C. gallina* is also affected by the concentration of the available food supply, as is the case for all filter-feeding organisms (Arneri et al., 1998; Häder et al., 2007). Chlorophyll concentration is a good food proxy for clams, based on the assumption that phytoplankton is the main component of suspension-feeding bivalves’ diet (Gillikin et al., 2005). Nowadays, the northern area of the western Adriatic basin is characterized by the presence of Po River run-off, which makes the NAS a eutrophic area with the highest average primary production and Chlorophyll *a* concentration in the Adriatic basin (Gilmartin et al., 1990; Mancuso et al., 2019). The higher growth rates of *C. gallina* found in present-day assemblages thus could also result from the current high food availability compared to past environmental settings of HCO, where, especially for early HCO, the geomorphologic setting of the study area was completely different. Indeed, wide estuaries with lagoon barrier systems developed along the targeted coastal areas (Amorosi et al., 2019). Thus, the modern study area shows a river-influenced delta where the freshwater and nutrient-rich plumes mix with nearshore marine waters. Early HCO settings were characterized by estuarine settings with barrier island/lagoon complexes, where freshwater input was mainly mixed within the estuaries and lagoons (Amorosi et al., 2019). Thus, sampled shoreface settings showed a reduced fluvial influence. Increasing temperatures will be associated with increased sea level in the coming decades (Lee et al., 2023; Vermeer and Rahmstorf, 2009), reducing the plumes of nutrient-rich freshwater in the Adriatic, with mixing occurring more within the river outlet.

Together with previous research, our study suggests that *C. gallina* can adapt to (limited) environmental changes resulting from human societal development (see also a recent review by Grazioli et al., 2022) and highlights its ability to persist in a scenario of limited global warming in the near future (e.g., < 1.5 °C change between the Holocene and modern-day settings in our study).

## 5. Conclusion

This study analyzes the variation in shell growth (i.e., linear extension and net calcification rates) of the bivalve *C. gallina* from assemblages from present-day and Holocene climate optimum when the SST in the investigated area of the Adriatic Sea was supposedly 1.5°C higher than today. The comparison between Holocene sub-fossil records and modern assemblages provided insights into the growth dynamics of this bivalve on a millennial temporal scale, overcoming the time limits imposed by laboratory studies. Shell growth decreased per unit of time with SST, showing reduced linear extension rate and calcification in warmer fossil horizons. Faster linear extension rates and higher calcification rates were found in present-day *C. gallina* populations.

The difference in growth dynamics across studies highlights how the combination of specific biotic and abiotic conditions at the local level can strongly influence the organism response. It also points to the high plasticity of this species when adapting to changing environments. Specifically, the present-day environmental context appears more favorable for *C. gallina* growth than the HCO. Given that the HCO may serve as an analog of near-future climate-environmental dynamics, we hypothesize that *C. gallina* will suffer a reduction in growth rate mainly due to climate warming and related environmental changes. Further investigation into the genetic variability of HCO *C. gallina* populations and in living populations along the latitudinal gradient in the Adriatic Sea could give insights into a selection of morphotypes more adapted to warmer waters and provide a better understanding of the degree of plasticity in *C. gallina* populations compared to local adaptation in the face of anthropogenic-derived climate warming.

## CRediT authorship contribution statement

**Alessandro Cheli**:. **Arianna Mancuso**:. **Fiorella Prada**:. **Alexis Rojas**: Writing – review & editing. **Giuseppe Falini**:. **Stefano Goffredo**:. **Daniele Scarponi**:.

## Data availability

All data generated or analyzed during this study are included in this published article and its online supplementary material.

## Acknowledgments

This study partially fulfills the requirements for a PhD thesis of A. Cheli, the PhD Course of Innovative Technologies and Sustainable Use of Mediterranean Sea Fishery and Biological Resources (FishMed-PhD) (University of Bologna, Italy). This work was supported by 1) the European Union–NextGenerationEU through the Italian Ministry of University and Research under Piano Nazionale di Ripresa e Resilienza (PNRR): Mission 4 Component C2, Investment 1.1 “Conservation of life on Earth: The fossil record as an unparalleled archive of ecological and evolutionary responses to past warming events”; 2) the National Recovery and Resilience Plan (NRRP), Mission 4 Component 2 Investment 1.4 - Call for tender No. 3138 of 16 December 2021, rectified by Decree n.3175 of 18 December 2021 of Italian Ministry of University and Research funded by the European Union – NextGenerationEU. Project code CN_00000033, Concession Decree No. 1034 of 17 June 2022 adopted by the Italian Ministry of University and Research, CUP J33C22001190001, Project title “National Biodiversity Future Center - NBFC”.

## Appendix A. Supplementary data

Supplementary material.

## Table captions

**Table 2. Mean value and standard error of shell skeletal and growth parameters.** Values for each assemblage in chronological and temperature order. K-W = Kruskal-Wallis rank test; NS not significant, * p < 0.05, *** p < 0.001. Assemblage labels, length (anterior-posterior maximum distance) and bulk density are after (Cheli et al., 2021).

