## Supplementary Material for "Insights on *Chamelea gallina* growth dynamics from the Holocene climate optimum in the Northern Adriatic Sea (Italy)"

### Online Supplementary Material
### Cheli et al.

#### Supplemental Methods

**Specimens' collection (from Cheli *et al.*, 2021)**

In particular, the two Holocene horizons were collected from core 205-S6 (Comacchio, 44°68'N, 12°15'E), respectively, at depths of 17.2 m (code "CO1") and 13.2 m (code "CO2"). Both horizons came from shoreface depositional environments characterized by sandy substrates and estimated water depths between 5 and 10 m.

Modern samples of *C. gallina* were collected in the Northern Adriatic Sea off the coast of Goro (code "MGO"; 44°75'N, 12°43'E) and Cervia (code "MCE"; 44°30'N, 12°40'E), two sites near to the extraction areas of cores used in this study and distant about fifty kilometers away from each other. Samplings were performed during scuba diving and using Van Veen Grab on the sandy bottom, ranging from 1 to 5 m water depth. Sampling operations were confined to the top-most 10 cm of the sea bottom's taphonomically active zone (TAZ). This sampling allowed us to collect a time-averaged record of shells estimated in tens of years, following the deposition rates reported in Trincardi *et al*. This allowed a better comparison between modern and sub-fossil horizons, consisting of shells from tens of years. No living organism was collected for this study.

Only 5–30 mm long shells (the maximum distance on the anterior-posterior axis) were considered for the analyses. The technical difficulties in obtaining reliable measurements in very small specimens defined the lower size limit. The upper size limit was due to the difficulty of collecting whole shells over 30 mm in the sub-fossil horizons with a 90 mm core diameter.

Before any measurements, each valve was cleaned with a toothbrush and soaked in distilled water for two hours to remove any external residue on the shell's surfaces. In addition, valves from modern settings were immersed in distilled water and hydrogen peroxide (5% vol.) for 24 h to eliminate any traces of organic material (e.g., epibionts). Then, the valves were dried in an oven at 37°C for one night to remove any moisture that may influence subsequent measurements.

**Environmental Parameters (from Cheli *et al.,* 2021)**

Sea surface temperature (SST) for the Adriatic Sea of modern settings was obtained from the global ocean OSTIA sea surface temperature and sea ice analysis databank (https://ghrsst-pp.metoffice.gov.uk). Mean annual SST was calculated from daily values from January 2010 to December 2019 (number of daily values = 3651 for each site). SST estimates for sub-fossil horizons were based on the Alkenones unsaturation index, a widely applied proxy for past SST and obtained from the high-resolution SST record of the past 10,000 years for the central-northern Mediterranean Sea (Gulf of Lion; Jalali *et al.,* 2016), considered a comparable physiographic setting for the North Adriatic Sea.

**Shell biometrics and skeletal parameters (from Cheli et al. 2021, slightly modified)**

Shell length (the maximum distance on the anterior-posterior axis) was measured using ImageJ software after data capture of each shell shape with a scanner (Acer Acerscan Prisa 620 ST 600 dpi), and dry shell weight was measured using an analytical balance (±0.1 mg).

Shells of different sizes from each assemblage were used for the analyses. They were divided into two groups according to size: immature shells (up to 18 mm) and mature ones (over 18 mm).

Shell skeletal parameters were measured by buoyant weight (BW) analysis using a density determination kit Ohaus Explorer Pro balance (± 0.1 mg; Ohaus Corp., Pine Brook, NJ, USA) as reported in Gizzi *et al*. 2016.

The BW measurement was repeated three times, and the average was considered for statistical analysis. The BW technique allowed us to estimate the variable of interest:

i. micro-density (mass per unit volume of the material which composes the shell, excluding the volume of pores; $g \cdot cm^{-3}$);
ii. apparent porosity: the volume of pores connected to the external surface (%);
iii. bulk density: shell mass/volume ratio, including the volume of pores, the volume of pores; $g \cdot cm^{-3}$).

At the microscale level, the shells were composed of pure aragonite with a perfectly preserved mineral phase and no relevant diagenetic alteration, except for a slight degradation of the inter-crystalline organic phase retrieved (Cheli *et al.,* 2021).

**Shell aging methods details**

For counting surface external rings, which appeared as smooth clefts on the shell surface and as strong pigmented lines across the anterior-posterior axis, shells were scanned in transmitted light to enhance the contrast of the surface's ridge and highlight bands at different densities.

To estimate age using shell sectioning, the valves were embedded in epoxy resin under vacuum at room temperature, followed by 24-hour hardening. The shells were sectioned along the anterior-posterior axis, from the umbo to the ventral margin, using an electrodeposited diamond cutting blade for counting internal bands. Sections were ground using successive finger grits (600, 1200, 2400 μm) and polished with abrasive alumina compound (3M Perfect-it III Extrafine Paste). Finally, the sections were ultrasonically cleaned, rinsed in purified water, and dried. Shell sections were then examined and photographed under oblique light at low magnification to identify internal growth bands. In each section, the number of annual growth rings was determined by counting the alternating opaque (carbonate matrix) and translucent (carbonate-organic matrix) increments visible on the shell cross-section (Arneri E et al., 1995) using a dissecting microscope under reflected light at low magnification (6.4 X). Assuming that the growth rings are laid down yearly, the age of each clam was estimated by counting all the translucent zones.

To validate the data from the two counting rings methods, oxygen isotopic measurements ($\delta^{18}O$) were carried out on "spot" samples collected sequentially from the umbo to the ventral edge of 3 shells of different dimensions for each site. Dried homogenized powdered samples were treated with helium, and then an acidified solution consisting of 104% orthophosphoric acid was added and left to react for 1 hour at 70 °C. Each sample was analyzed using a Thermo Gasbench preparation system attached to a Thermo Delta V Advantage mass spectrometer in continuous flow mode. Each run of samples was accompanied by ten reference carbonates (Carrara Z) and two control samples (Fletton Clay). Carrara Z has been calibrated to VPDB using the international standard NBS19.

#### Supplemental Figures

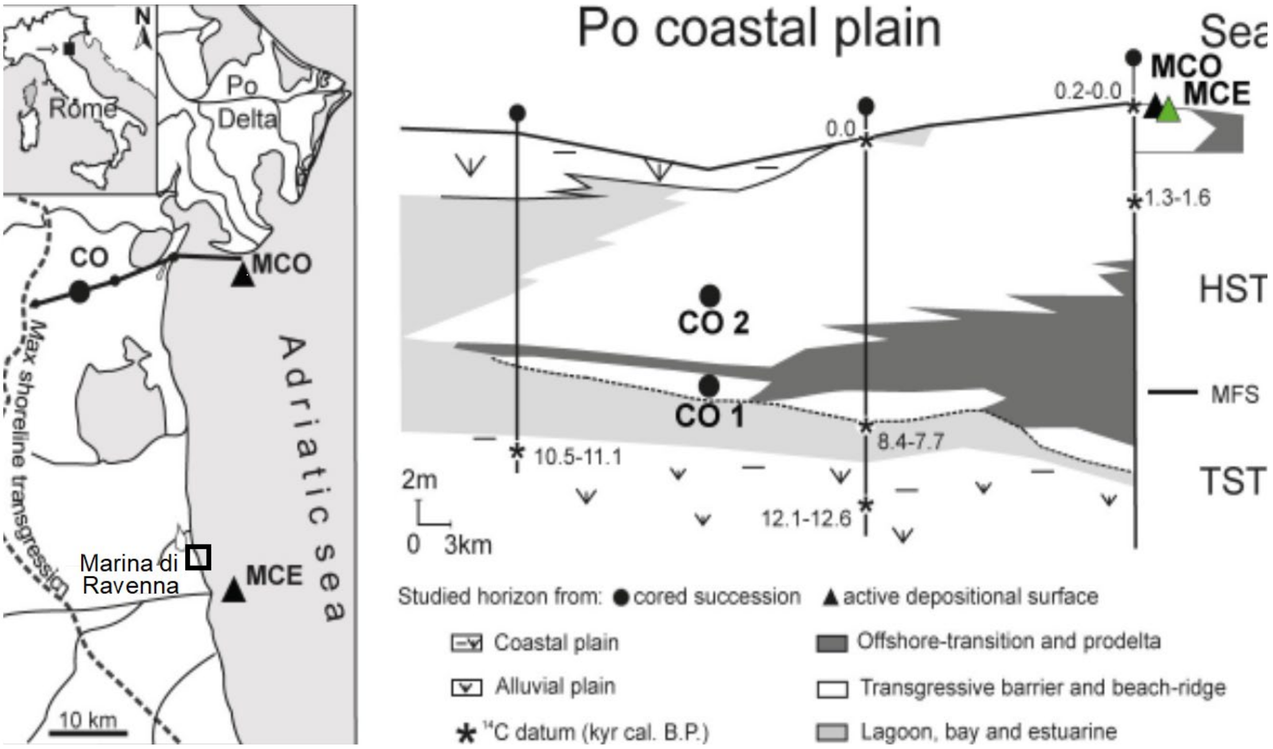

**Figure. S1**. Study area map and stratigraphic framework of the latest Quaternary Po coastal plain-sedimentary succession (revised by Cheli *et al.,* 2021). In the left panel, black dots mark HCO assemblages (CO1, CO2), black triangles mark sampled thanatocoenosis from present-day shoreface environments (MCE, MCO), square marks the location of Marina di Ravenna, and the black solid line represents the along-dip cross-section of the Po coastal plain (right panel). The right panel shows sketched climate-driven environmental changes within the study area during the Holocene; the dashed line indicates the wave ravinement surface and green-filled symbols represent projected samples on the cross-section. Acronyms: MFS = maximum flooding surface; TST = transgressive systems tract; HST = highstand systems tract.

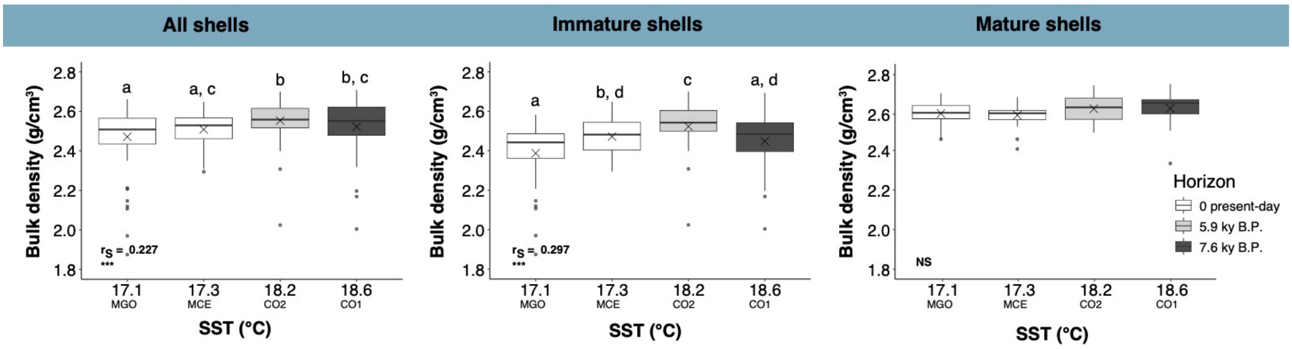

**Figure S2.** Variation of bulk density in relation to different SST (Sea Surface Temperature). Three groups were considered: correlations in all shells, correlations in immature shells (<18 mm), and correlations in mature shells (>18 mm). The box colors represent the age of the assemblages, ranging from the oldest fossil horizon (dark grey boxes) to the present-day horizon (white boxes). The boxes indicate the 25$^{th}$ and 75$^{th}$ percentiles, the lines within the boxes mark the medians, and the crosses mark the mean values. Black points represent outliers. Different letters indicate statistical differences among assemblages ($p < 0.05$; the number of clams measured for each assemblage is reported in Table 1). $r_S$, Spearman's rho coefficient. ***$p < 0.001$, NS, not significant.

### Supplemental Table

**Table S1**. Radiocarbon data for individual specimens of *Chamalea gallina* from core 205-S6 sampled at 17.10 m core depth and used for estimation of time-averaging of sample CO1 (i.e., interquartile range).

| SAMPLE DESCRIPTORS | | | | 14C DESCRIPTORS AND RADIOMETRIC AGE | | | | | | CALIBRATED AGE (Y.B.P.) | | |
|---|---|---|---|---|---|---|---|---|---|---|---|---|
| **Core ID** | **Sample ID** | **depth (m)** | **Facies** | **Fraction modern** | **error** | **D 14C** | **error** | **Age 14C** | **error** | **median** | **2s old** | **2s yng** |
| 205-S6 | CO1-1 | 17.10 | shoreface | 0.400426571 | 0.003118 | -599.573 | 3.117556 | 7350 | 70 | 7679 | 7829 | 7556 |
| 205-S6 | CO1-2 | 17.10 | shoreface | 0.41499394 | 0.003169 | -585.006 | 3.168601 | 7060 | 70 | 7435 | 7559 | 7304 |
| 205-S6 | CO1-3 | 17.10 | shoreface | 0.402457845 | 0.00322 | -597.542 | 3.219648 | 7310 | 70 | 7640 | 7789 | 7507 |
| 205-S6 | CO1-4 | 17.10 | shoreface | 0.40290253 | 0.002906 | -597.097 | 2.905615 | 7300 | 60 | 7632 | 7771 | 7510 |
| 205-S6 | CO1-5 | 17.10 | shoreface | 0.404109428 | 0.003101 | -595.891 | 3.100696 | 7280 | 70 | 7612 | 7754 | 7481 |
| 205-S6 | CO1-6 | 17.10 | shoreface | 0.408723272 | 0.002965 | -591.277 | 2.964883 | 7190 | 60 | 7532 | 7647 | 7421 |
| 205-S6 | CO1-7 | 17.10 | shoreface | 0.403931637 | 0.003427 | -596.068 | 3.426882 | 7280 | 70 | 7616 | 7770 | 7478 |
| 205-S6 | CO1-8 | 17.10 | shoreface | 0.405556803 | 0.003716 | -594.443 | 3.71624 | 7250 | 80 | 7586 | 7736 | 7437 |
| 205-S6 | CO1-9 | 17.10 | shoreface | 0.415306016 | 0.003745 | -584.694 | 3.745432 | 7060 | 80 | 7429 | 7563 | 7282 |
| 205-S6 | CO1-10 | 17.10 | shoreface | 0.406444916 | 0.003029 | -593.555 | 3.02942 | 7230 | 60 | 7571 | 7688 | 7443 |
| 205-S6 | CO1-11 | 17.10 | shoreface | 0.40195615 | 0.001067 | -601.4 | 1.067254 | 7320 | 25 | 7641 | 7726 | 7568 |
